# Hypoxia exposure fine-tunes mitochondrial function in sea turtle cells

**DOI:** 10.1101/2024.11.01.621592

**Authors:** B. Gabriela Arango, David C. Ensminger, Dianna Xing, Celine A. Godard-Codding, José Pablo Vázquez-Medina

## Abstract

Sea turtles experience extreme fluctuations in oxygen availability derived from extended breath-hold diving, yet the cellular adjustments that underlie hypoxia tolerance in these animals remain poorly understood. Here, we used metabolite profiling, extracellular flux assays, and microscopic analyses of the mitochondrial reticulum to study how primary cells derived from sea turtles and lizards cope with extended hypoxia exposure. Cells from both species proliferate in primary culture, stain positive for fibroblast markers, are metabolically active, and stabilize HIF1-α when exposed to chemical or environmental hypoxia. In contrast to lizard cells, sea turtle cells exhibit a faster and more robust response to 1 or 24-hour hypoxia exposure (0.1% O_2_), upregulating antioxidant pathways and optimizing oxygen use rather than relying on glycolytic metabolism. Similarly, the mitochondrial reticulum is maintained without apparent fragmentation during hypoxia exposure in sea turtle cells. Consistent with these observations, mitochondrial function readily recovers in sea turtle but not lizard cells upon reoxygenation. These findings show that sea turtle cells undergo intrinsic metabolic adjustments to cope with extreme oxygen fluctuations, aligning with the remarkable hypoxic tolerance exhibited by these animals, which can endure up to 7 hours of breath-holding underwater.

## INTRODUCTION

Hypoxia-inducible factors (HIFs) orchestrate the canonical response to low oxygen tension by regulating genes involved in oxygen use, glycolytic energy production, and cell survival (Wang et al., 1995; Kim et al., 2006; Semenza, 2014). Whereas this response was first described in mammalian cells, it appears to be conserved across the animal kingdom (Wenger, 2002; Rytkönen et al., 2007, 2011; Gorr et al., 2010; Majmundar et al., 2010; Graham and Presnell, 2017; Zhu et al., 2018; Lee et al., 2020). Shifting intermediary metabolism from oxidative phosphorylation to glycolysis alters mitochondrial fission and fusion dynamics, resulting in structural changes in the mitochondrial reticulum (Chandel and Schumacker, 2000; Lee and Finkel, 2013; Kumar and Choi, 2015; Lee et al., 2020, 2021). Furthermore, biochemical events during hypoxia exposure, such as succinate accumulation from reverse electron transfer between mitochondrial complex I and complex II and conversion of xanthine dehydrogenase into xanthine oxidase, increase oxidant generation upon reoxygenation, potentially compromising mitochondrial function and cell survival (Brown et al., 1988; Chouchani et al., 2016; Sies and Jones, 2020).

Animals adapted to natural changes in oxygen availability employ diverse strategies to adjust metabolism and cope with reoxygenation-induced oxidant generation, such as upregulating antioxidant defenses during hypoxia (Hermes-Lima and Zenteno-Savín, 2002; Zhou et al., 2018). In anoxia-tolerant freshwater turtles, HIF-1 promotes metabolic rewiring of oxygen utilization (Milton and Prentice, 2007; Sparks et al., 2022), which likely contributes to maintaining mitochondrial integrity (Bundgaard et al., 2019b) and restarting mitochondrial respiration upon reoxygenation (Fago, 2022). Similarly, freshwater turtle tissues suppress mitochondrial complex V activity (Gomez and Richards, 2018) and avoid succinate accumulation during hypoxia (Bundgaard et al., 2019a), preventing oxidant generation during reoxygenation. Sea turtles spend hours underwater (Hochscheid et al., 2005), but it is unknown if similar metabolic adjustments allow them to withstand extended hypoxia exposure.

Previous work suggests that cells derived from sea turtles stabilize HIF-1 faster than human cells upon hypoxia exposure (Barlian and Riani, 2018), but comparative cellular analyses of diving and non-diving reptiles are lacking. Here, we studied the response to hypoxia in primary cells derived from sea turtles using a comparative approach. Our results show that while hypoxia exposure stabilizes HIF-1 in both sea turtle and lizard cells, sea turtle cells cope with hypoxia exposure by upregulating antioxidants and preserving the mitochondrial reticulum, which promotes efficient recovery of cellular respiration upon reoxygenation.

## METHODS

### Animals and tissue collection

Animal work was approved by San Jose State University’s IACUC #1078. Female Western fence lizards (*Sceloporus occidentalis*) were collected in the vicinity of the University of California Santa Cruz, under permit CDFW S-212470001-21248-001, and transported to San Jose State University for separate experiments. Hatchlings were later euthanized as juveniles following our previously published protocols (Ensminger et al., 2018). Tail and dorsal skin tissue samples were rinsed with 70% ethanol and PBS (Gibco #14190-144), placed in complete fibroblast growth medium consisting of DMEM/F-12 (Gibco #10565018) supplemented with 10% fetal bovine serum (FBS, Seradigm Life Science #89510-186), 1% penicillin-streptomycin (Gibco #15140-122) and 1% 1M HEPES (Gibco #15630-056), and transported on ice to the University of California, Berkeley. Primary cells were isolated within 3 hours of tissue collection. Loggerhead sea turtle (*Caretta caretta*) cells were derived from the skin, as described previously by Webb et al., 2014.

### Isolation and culture of primary cells

Lizard tissues were minced in DMEM/F-12 without FBS, transferred to a conical tube containing 0.67 mg/ml Collagenase II (Worthington #LS004176) diluted in DMEM/F-12, and placed in a CO_2_ incubator at 26°C for 24 hours. Samples were centrifuged, and the pellet was rinsed twice with PBS, resuspended in complete growth media, seeded in gelatin-coated tissue culture flasks, and incubated overnight. Complete growth media was added to each flask two days after initial cell plating, and the media was replaced a day later. Lizard cells were maintained at 26°C with 5% CO_2_.

Confluent cells were dissociated with 0.05% Trypsin-EDTA (Gibco #25300-054), expanded, and cryopreserved in liquid nitrogen at P4. Previously established sea turtle dermal fibroblasts were grown at 28°C with 5% CO_2_. The media consisted of RPMI-1640 (Thermo Fisher Scientific #11835030) supplemented with 10% FBS (Seradigm Life Science), 1% penicillin-streptomycin (Gibco #15140-122), 1% Amphotericin B (Thermo Fisher Scientific #15290018) and 100 mM HEPES (Gibco). Confluent sea turtle cells were dissociated with 0.25% Trypsin-EDTA (Gibco #25200-056), expanded, and cryopreserved in liquid nitrogen.

### Cell characterization

We generated cell lines from five individuals per species. Pooled stock cultures from similar passages were created from all individuals and used for experiments. The day before the experiments, all cells were switched to the DMEM/F-12 media formulation and similar temperature conditions. The fibroblast-like phenotype of the cells was confirmed by immunofluorescence for vimentin and platelet-derived growth factor receptor (PDGFRα).

### Immunofluorescence

Immunofluorescence was conducted as previously described (Vázquez-Medina et al., 2016). Briefly, 50,000 cells were grown in gelatin-coated 35mm glass-bottom dishes, fixed with 1:1 ice-cold acetone/methanol, permeabilized, blocked, and incubated overnight with antibodies against vimentin (Cell Signaling Technology #5741S diluted 1:100), platelet-derived growth factor receptor α (PDGFRα, Cell Signaling Technology #3164S diluted 1:200), or hypoxia-inducible factor 1α (HIF-1α, Cell Signaling Technology #36169 diluted 1:100). Alexa fluor plus 488 or 594 secondary antibodies were used at 1:200 dilution. Nuclei were counterstained with NucBlue (Invitrogen #R37606). Cells were imaged using a Zeiss Axio Observer inverted fluorescence microscope fitted with a 63X objective and Zen software. Negative controls with no primary antibody were included for each antibody and cell type.

### Hypoxia exposure

Chemical hypoxia was induced by overnight incubation with 250 μM or 500 μM CoCl_2_, a hypoxia-mimetic agent that stabilizes HIF-1α (Muñoz-Sánchez and Chánez-Cárdenas, 2019). In separate experiments, cells were exposed to hypoxia using an InVivO_2_ physiological cell culture workstation (Baker Ruskinn) as described previously (Allen et al., 2024). Hypoxic cell culture conditions were set at 0.1% O_2_, 5% CO_2_, and 26°C for 1 or 24 hours. Control conditions (normoxia) were 21% O_2_, 5% CO_2_, and 26°C. For hypoxia/reoxygenation experiments, cells were incubated for 1 hour at 0.1% O_2_ followed by 1-hour reoxygenation or 24 hours at 0.1% O_2_ followed by 1-hour reoxygenation. We used live/death cell assays (Invitrogen #L34973) to confirm that cells from both species remained viable after exposure to 0.1% O_2_ for 24 hours. Cells collected within the hypoxic environment or after hypoxia/reoxygenation were used for HIF-1α staining, metabolite profiling, mitochondrial respiration, or mitochondrial network analyses.

### Metabolite profiling

Hypoxic cells were flash-frozen in liquid nitrogen. Primary carbon metabolite profiling was conducted at the UC Davis West Coast Metabolomics Center using an ALEX-CIS gas chromatography-mass spectrometry (GC-TOF MS) platform, following the methods of Fiehn et al., 2008. Each sample was run in triplicate. Raw data were pre-processed and stored as apex masses, exported to a data server with absolute spectra intensities, and further filtered with an algorithm implemented in the BinBased database (Fiehn et al., 2008). Spectra were cut to a 5% base peak abundance and matched to a database entry. Quantification was stored as peak height using the unique ion as default for all database entries that are positively detected in more than 10% of the unidentified metabolites, as described in our previous work (Arango et al., 2022). We analyzed peak intensities with a 5% variance filter using the interquartile range for untargeted metabolomics and a 5% abundance filtering for minimal values using the mean. Data were normalized to the median.

### Extracellular flux assays

Oxygen consumption rates (OCR) were measured at 26°C using a calibrated Seahorse XFp Extracellular Flux Analyzer (Agilent Technologies, MA). Cells were seeded in gelatin-coated Seahorse mini plates (Agilent Technologies). Seeding density and carbonyl cyanide-4 (trifluoromethoxy) phenylhydrazone (FCCP) concentrations were optimized experimentally for each species. The optimal cell density was 40,000 cells/well for lizard cells and 20,000 cells/well for sea turtle cells. On the day of the assay, cells (n=6) were washed with Seahorse XF DMEM assay medium (pH 7.4), supplemented with 10 mM glucose (Agilent #103577-100), 1 mM pyruvate (Agilent #103578-100) and 2 mM L-Glutamine (Agilent #103579-100), and incubated at 26°C in a CO_2_-free incubator for 1 hour. OCR was measured using the Cell Mito Stress Test kit (Agilent #103015-100) at 26°C, with oligomycin (1.5 μM), FCCP (0.4 μM for lizard cells and 2 μM for sea turtle cells), and 0.5 μM rotenone/antimycin A. OCR values were normalized to cell count and total protein content measured using a Qubit protein assay (Invitrogen #Q33211). Basal respiration was calculated as oxygen consumption before any drugs were added, minus the non-mitochondrial oxygen consumption, measured using rotenone (complex I inhibitor) and antimycin A (complex III inhibitor). We used oligomycin to calculate respiration driving mitochondrial ATP synthesis, whereas proton leak was calculated as protons across the inner membrane, independently of ATP synthesis. Maximal respiration was calculated after injecting FCCP. Spare respiratory capacity was calculated using maximal and basal respiration (Divakaruni et al., 2014).

### Mitochondrial network analyses

50,000 cells seeded in gelatin-coated, glass-bottom dishes were stained with 100 nM MitoTracker Red-CMXRos (Cell Signaling Technology #9082) and NucBlue (Hoescht 33342, Invitrogen) and exposed to hypoxia or hypoxia/reoxygenation. Cells were fixed with ice-cold methanol, as previously described (Torres-Velarde et al., 2021) and imaged using a Zeiss Axio Observer fluorescence microscope fitted with a 63X objective and Zen software. Fifty cells were randomly selected from each of the three replicates per treatment. Mitochondrial footprint, branch length, and mitochondrial networks were quantified from binary raw images using the MiNa (v3.0.1) FIJI plugin (Valente et al., 2017).

### Statistical Analyses

Statistical analyses and visualization were conducted in R Studio (R Core Team, 2022; Gearty and Jones, 2023). Groups were compared using two-sample *t*-tests or ANOVA. Normal distribution was assessed using Shapiro-Wilk tests. Equality of variances was assessed using Levene’s or Fisher’s F-test. Non-parametric data were log-transformed to meet model assumptions. Mitochondrial network comparisons were conducted using Kruskal-Wallis tests followed by pairwise Wilcox tests using the Benjamini-Hochberg method for p-value adjustment. Metabolite peak intensities were compared using multiple t-tests with an FDR correction of 5%. Statistical analyses were conducted using Metaboanalyst 6.0 (Xia et al., 2009; Main server: https://www.metaboanalyst.ca, 14 October 2024) and R studio. Statistical significance was considered at p ≤ 0.05. Differentially abundant metabolites were called based on adjusted p values and Log2FCs of ± 1. Enrichment analyses were conducted using the Small Molecule Pathway Database (SMPDB), with sets containing at least three entries using the global test method (Goeman et al., 2004).

## RESULTS

### Reptile fibroblasts show species-specific bioenergetic profiles

We derived primary dermal fibroblasts from western fence lizards and loggerhead sea turtles to study the cell-autonomous response to low oxygen (hypoxia) and hypoxia/reoxygenation in diving and non-diving reptiles. Cells from both species proliferate in primary culture and stain positive for the fibroblast markers PDGFRα and vimentin (**Figure 1a**). Similarly, cells from both species respire and respond to oligomycin, FCCP, and rotenone/antimycin A (**Figure 1b**). Basal, maximal, spare, and ATP-linked respiration were significantly lower in lizard than in sea turtle cells during normoxic conditions (Basal: 10.6 ± 2.20 vs. 19.8 ± 2.09 pmol/min/ug/mL, t = -7.42, df = 10, n = 12, p = 0.000023; Maximal: 25.9 ± 6.49 vs. 46.4 ± 3.82 pmol/min/ug/mL, t = -6.65, df = 10, n = 12, p = 0.000057; Spare: 15.3 ± 4.30 vs. 26.6 ± 2.32 pmol/min/ug/mL, t = -5.63, df = 10, n = 12, p = 0.00022; ATP-linked respiration: 7.48 ± 1.5 vs. 17.5 ± 1.34 pmol/min/ug/mL, t = -12.18, df = 10, n = 12, p = 0.00000025; **Figure 1c**). These results show that dermal fibroblasts derived from sea turtles and lizards are viable and metabolically active in primary culture. Furthermore, these results suggest species-specific differences in mitochondrial function and potentially oxygen management are retained at the cellular level.

**Figure 1.**
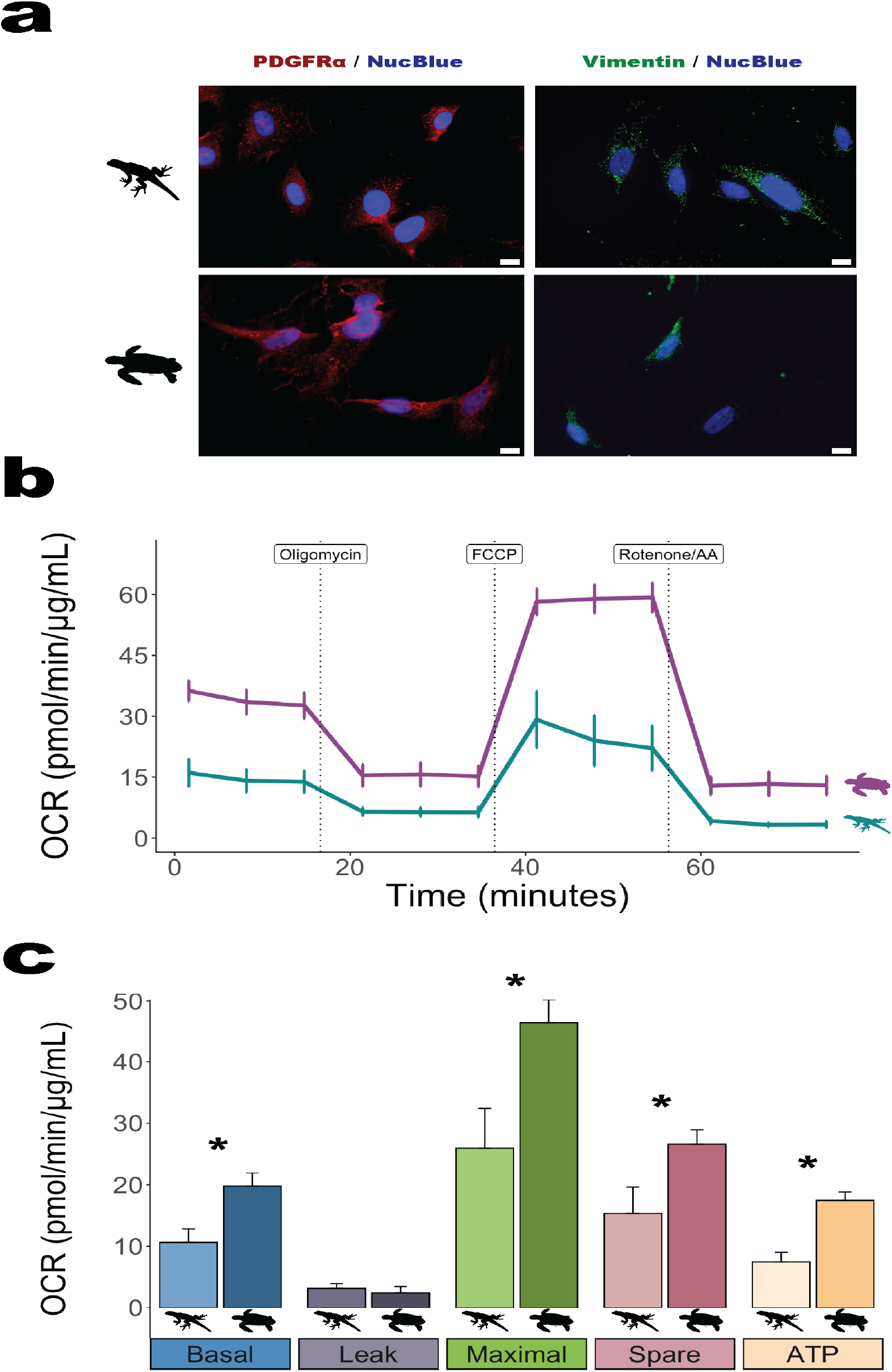
Reptile fibroblasts retain species-specific bioenergetic profiles in primary culture. (a) Immunofluorescence for PDGFRα and vimentin. (b) Cellular respiration in lizard and sea turtle cells measured using extracellular flux assays under normoxia. (c) Mitochondrial function: basal, proton leak, maximal, spare, and ATP-linked respiratory capacity, calculated according to Divakaruni et al., 2014. Stars indicate significant differences between species (p < 0.05, n=6).

### Hypoxia exposure stabilizes HIF-1α in reptile cells

We tested whether sea turtle and lizard cells respond to hypoxia by stabilizing HIF-1α. Both sea turtle and lizard cells stabilized HIF-1α when treated with 250 μM or 500 μM CoCl_2_, a chemical HIF-1α inductor (Muñoz-Sánchez and Chánez-Cárdenas, 2019) (**Figure 2**). These results show that reptile cells respond to chemical hypoxia by stabilizing HIF-1α and that the mammalian antibody reacts with reptile HIF-1α, likely due to the highly conserved nature of the protein (Graham and Presnell, 2017).

**Figure 2.**
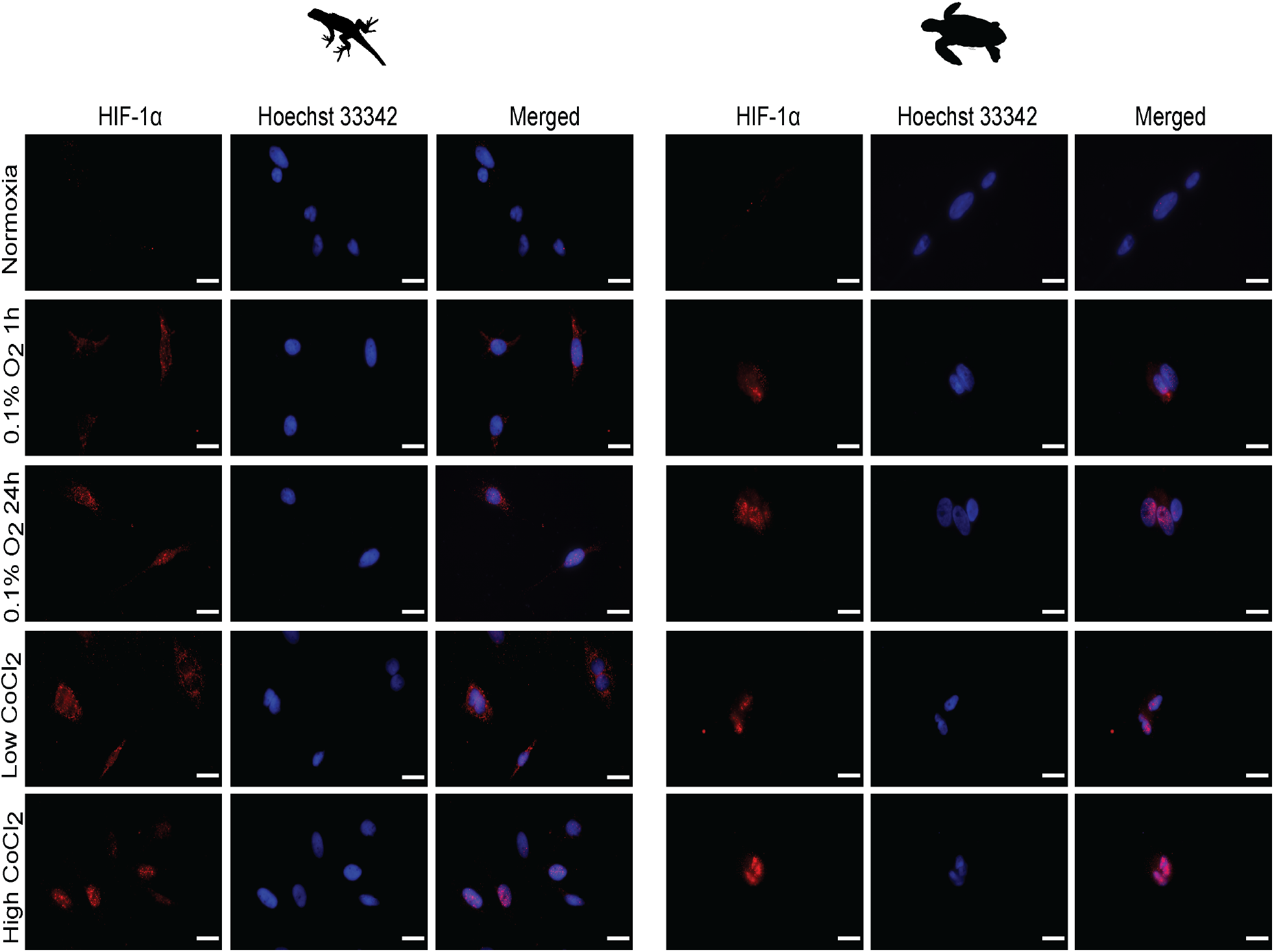
Hypoxia exposure stabilizes HIF-1α in reptile cells. Immunofluorescence for HIF-1α in lizard and sea turtle cells exposed to normoxia or 0.1% oxygen for 1 and 24h or treated overnight with 250 μM or 500 μM CoCl_2_.

Hypoxia exposure for 1h and 24h also resulted in HIF-1α stabilization in sea turtle and lizard cells, suggesting a conserved cellular response to low oxygen tension in reptiles. Of note, treatment with 250 μM CoCl_2_ or 1h hypoxia resulted in HIF-1α staining mainly in the perinuclear region in lizard cells, whereas treatment with 500 μM CoCl_2_ promoted a prominent nuclear HIF-1α staining. In contrast, all treatment conditions resulted in robust nuclear HIF-1α staining in sea turtle cells (**Figure 2**), suggesting potentially faster HIF-1α stabilization in diving than non-diving reptiles, as observed in primary cells derived from diving compared to non-diving mammals (Allen et al., 2024), and in sea turtle cells compared to human cells (Barlian and Riani, 2018).

### Sea turtle cells show distinctive metabolic signatures and upregulate antioxidants during hypoxia exposure

We conducted metabolite profiling to explore potential species-specific changes in primary carbon metabolites during hypoxia exposure in cells derived from diving and non-diving reptiles. Our analysis identified 1,023 metabolite peaks, of which 236 corresponded to known metabolites. Unsupervised hierarchical clustering showed significant differences in known metabolite abundance in response to hypoxia exposure between lizard and sea turtle cells (**Figure 3a**). Whereas the overall response to hypoxia, independent of time, resulted in a higher number of significantly different metabolites in sea turtle (66) compared to lizard cells (50) (**Figure 3b**), we detected opposite, species-specific patterns in response to 1h and 24h hypoxia exposure. After 1h hypoxia exposure, only 14 metabolites were significantly different in lizard cells, whereas 51 metabolites were significantly different in sea turtle cells. Interestingly, the abundance patterns flipped after 24h hypoxia exposure; the abundance of 36 metabolites was significantly different in lizard cells, but only 30 metabolites were significantly different in sea turtle cells. Of note, only 15 differentially abundant metabolites were shared between species when time was not considered, whereas 7 and 16 metabolites were shared at 1h and 24h hypoxia exposure, respectively (**Figure 3b**). These results suggest species-specific metabolic regulation in sea turtle and lizard cells with a faster and more robust response to hypoxia in sea turtle cells (**Figure 3b**). In lizard cells, the highest number of differentially abundant metabolites was observed after the 24-hour hypoxia exposure, where 14 metabolites were upregulated and 22 downregulated. In contrast, 19 metabolites were upregulated and 32 downregulated in sea turtle cells after only 1 hour of hypoxia exposure (**Figure 3c**).

**Figure 3.**
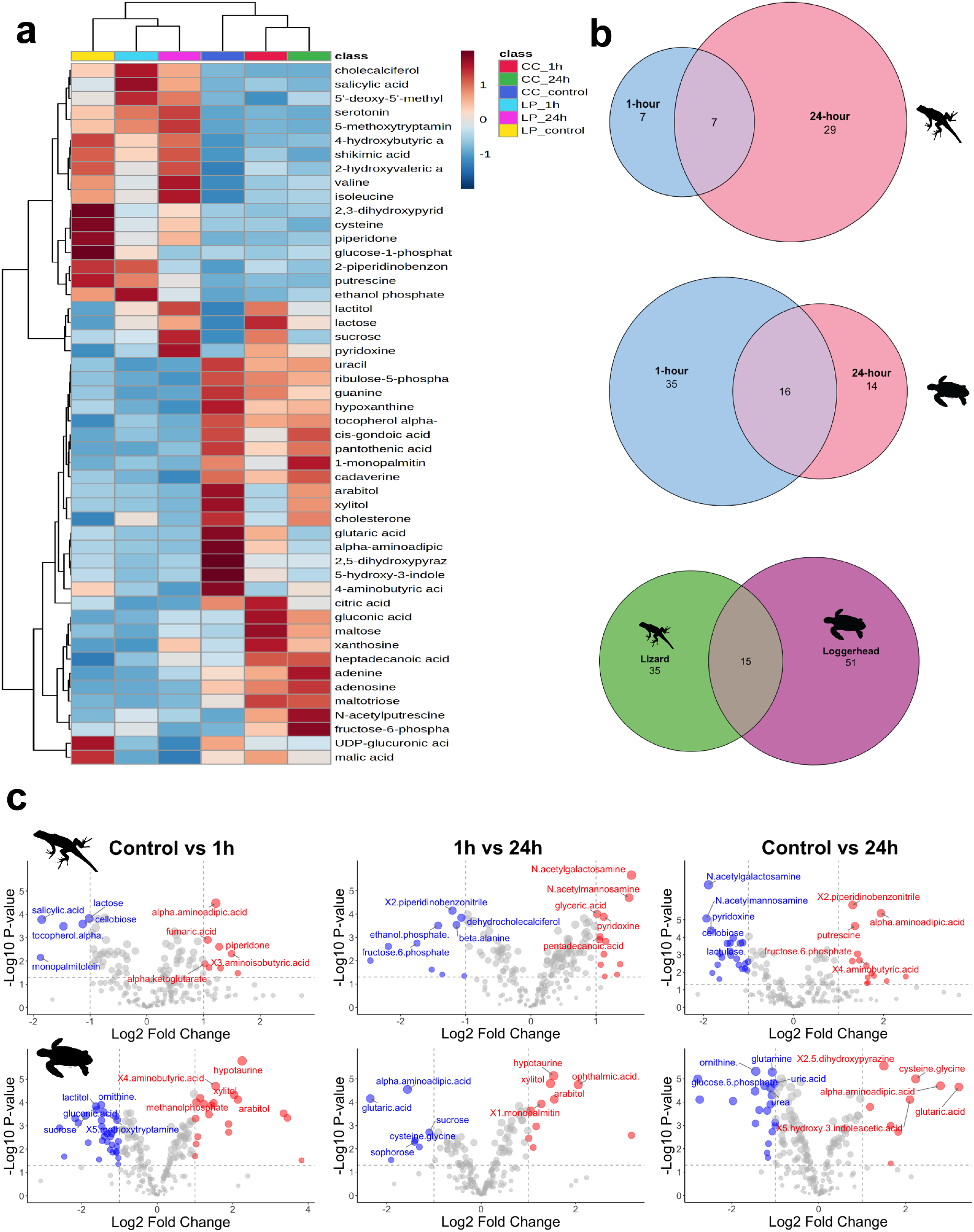
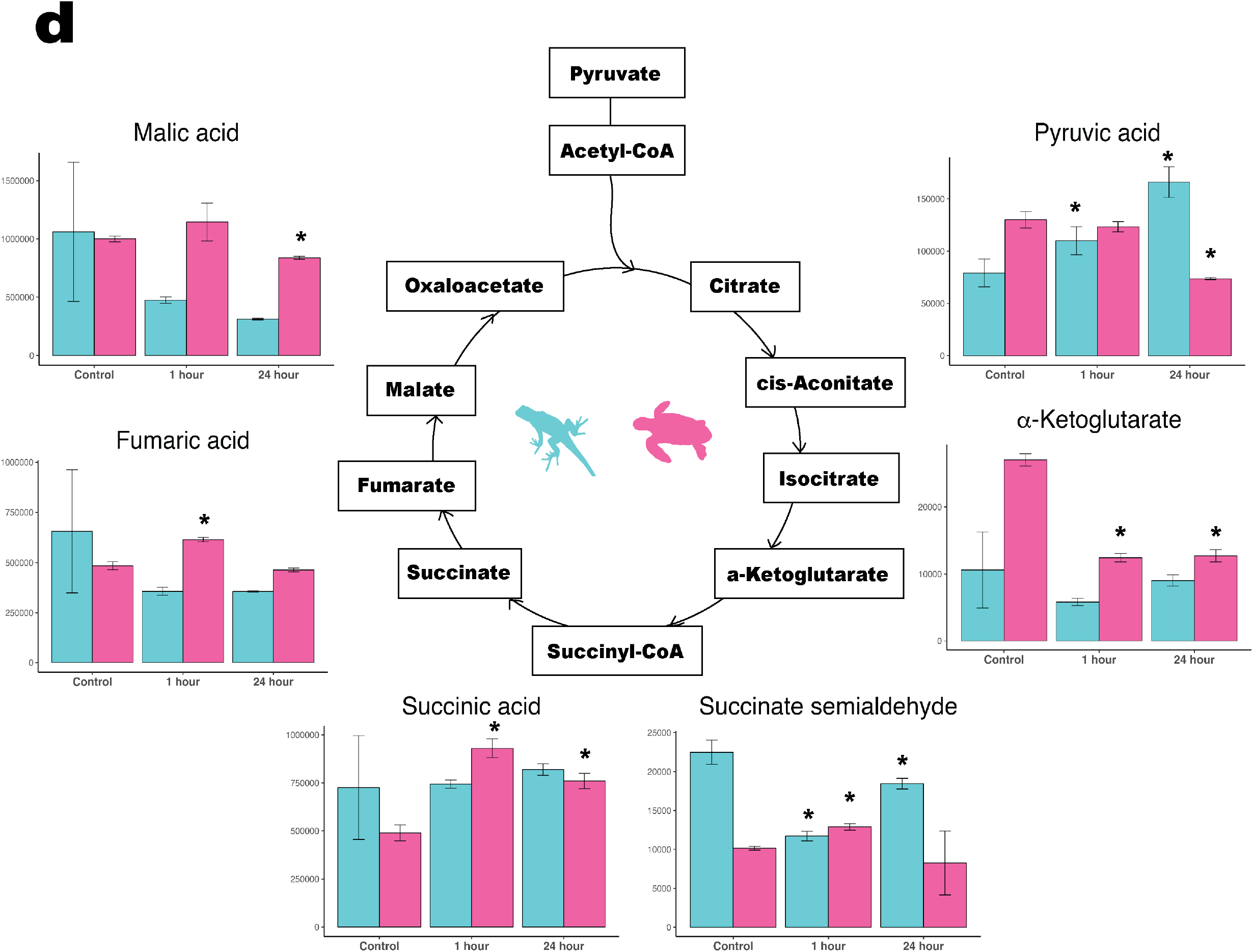
Sea turtle cells exhibit distinctive metabolomic signatures upon hypoxia exposure. (a) Hierarchical clustering showing the top 50 significantly abundant metabolites in lizard and sea turtle cells. (b) Venn diagrams comparing significantly different metabolites in lizard and sea turtle cells exposed to hypoxia. (c) Volcano plots highlight the top 10 regulated metabolites for each treatment. (d) Species-specific changes in TCA cycle metabolites. Peak heights represent the quantification ion (m/z value) at the specific retention index. One million hypoxic cells were used in each treatment. Stars denote significant differences between treatments and control for each species (p < 0.05).

We then conducted enrichment analyses with differentially abundant metabolites to detect relevant pathways that drive the response to short and long-term hypoxia exposure in reptile cells (**Supplementary Figure 1**). Metabolism of galactose, a glycolytic precursor that yields glucose, lactose, and other carbohydrate intermediaries, was one of the enriched pathways in lizard cells after 24-hour hypoxia exposure, with intermediates in this pathway downregulated. This suggests a reliance on glycolysis during oxygen-limited conditions. Consistent with this observation, glucose-1-phosphate, a glycolytic precursor and an end-product of galactose metabolism, was upregulated (**Figure 3d**). This response to hypoxia in lizard cells exemplifies the Crabtree effect, a short-term adaptation to hypoxia and substrate limitation that suppresses oxidative phosphorylation while using glycolysis to support ATP generation (Mookerjee et al., 2017).

In contrast to this observation, glucose metabolites were downregulated in sea turtle cells after 24h hypoxia exposure. Glycolysis receives inputs from at least seven metabolic pathways and outputs to three, including pyruvate metabolism. Sea turtle cells likely use an alternative pathway such as galactose or pyruvate metabolism (**Figure 3d**), thus circumventing the Crabtree effect. This enables continued use of oxidative phosphorylation for ATP generation in hypoxia. Furthermore, this poses a strategy to prevent the deleterious effects of reverse electron transport from decreased ATP production which can trigger cell death, as well as from reoxygenation damage via succinate accumulation (Eguchi et al., 1997; Chinopoulos, 2013).

Sea turtle cells also upregulated antioxidant metabolism, using glutamate as a precursor for glutathione, a major endogenous antioxidant (Willmore and Storey, 1997; Forman et al., 2009). Arginine, proline, and glutamate metabolism pathways were enriched in sea turtle cells after 1h hypoxia exposure. Proline, alanine, and glutamate are essential for the cellular stress response (Phang et al., 2010). During low oxygen conditions, proline catabolism indirectly supports ATP production by donating electrons to FADH2, a complex II substrate (Pallag et al., 2022). While increased complex II (CII) activity could result in reverse electron transport, the significant depletion of α-ketoglutarate at 1 hr and 24 hr in hypoxia combined with increased succinate indicates that electrons are moving in the forward direction through CII (**Figure 3d**).

This decreases the likelihood of oxidative damage from CII during reoxygenation in sea turtle cells. Hence, these results suggest that sea turtle cells mount an anticipatory response to oxidant stress when exposed to low oxygen tension, likely preparing to cope with reoxygenation-induced oxidant generation (Giraud-Billoud et al., 2019).

### Hypoxia exposure fine-tunes mitochondrial function in sea turtle cells

We conducted extracellular flux assays to study how sea turtle cells recover after hypoxia/reoxygenation and whether the observed species-specific differences in primary carbon metabolites relate to functional changes in cellular respiration. Exposure to short-(1h) and long-term (24h) hypoxia/reoxygenation elicited different responses in sea turtle and lizard cell respiration (**Figure 4a-4b**). Notably, sea turtle cells exhibit a steady downregulation of cellular respiration with increasing hypoxia exposure time (**Figure 4c**) and show faster recovery after extended hypoxia (24h) (**Figure 4d)** compared to lizard cells. While maximal respiration and spare respiratory capacity were lower in sea turtle than in lizard cells after 1h hypoxia/reoxygenation (Maximal: 35.7 ± 5.16 vs. 20.4 ± 2.6 pmol/min/ug/mL, t = 4.6, df = 4, n = 6, p = 0.01; Spare: 20.9 ± 2.31 vs. 8.58 ± 1.45 pmol/min/ug/mL, t = 7.8, df = 4, n = 6, p = 0.001; **Figure 4e**), basal, maximal and ATP-linked respiratory capacity were significantly higher in sea turtle compared to lizard cells after 24h hypoxia/reoxygenation (Basal: 4.37 ± 0.4 vs. 5.74 ± 0.33 pmol/min/ug/mL, t = -4.56, df = 4, n = 6, p = 0.01; Maximal: 8.15 ± 0.86 vs. 10.5 ± 1.12 pmol/min/ug/mL, t = -2.86, df = 4, n = 6, p = 0.04; ATP-linked: 2.7 ± 0.39 vs. 5.10 ± 0.32 pmol/min/ug/mL, t = -8.17, df = 4, n = 6, p = 0.001; **Figure 4f**) whereas proton leak was lower (Proton leak: 1.66 ± 0.03 vs. 0.65 ± 0.08 pmol/min/ug/mL, t = 19, df = 4, n = 6, p = 0.00004; **Figure 4f**). Although decreased mitochondrial function during reoxygenation was observed in sea turtle cells, reoxygenation damage is far more evident in lizard cells. Specifically, the response to FCCP, a proton shuttling uncoupler, is not sustained in Lizard cells during reoxygenation, suggesting damage to mitochondrial complexes, decreased proton pumping across the membrane, or a compromised mitochondrial membrane which would be unable to retain a large proton gradient. These results demonstrate species-specific regulation of cellular respiration and mitochondrial function after hypoxia/reoxygenation exposure. Furthermore, these results show that sea turtle cells are intrinsically better equipped to restart mitochondrial function after hypoxia exposure.

**Figure 4.**
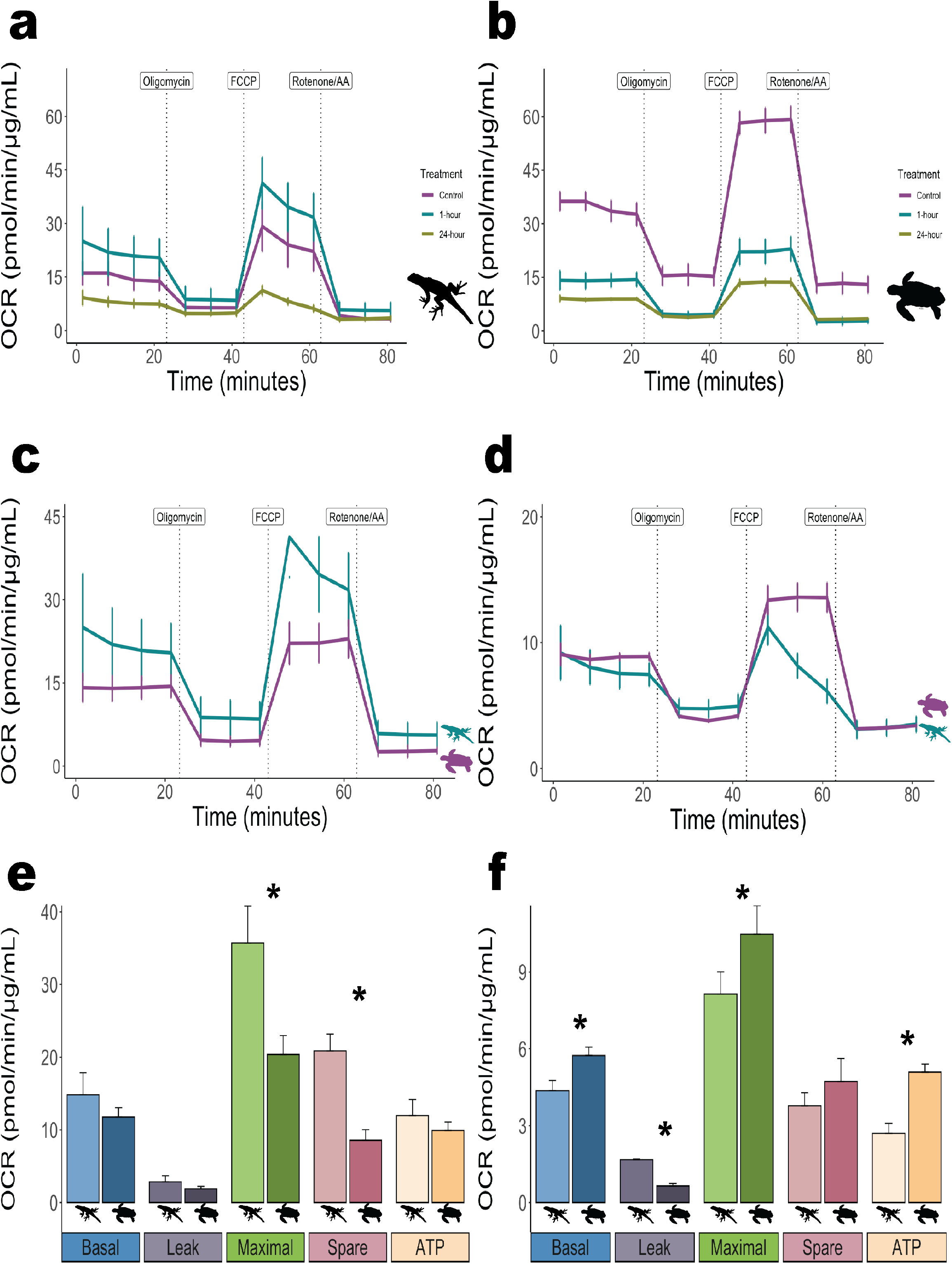
Hypoxia exposure fine-tunes mitochondrial function in sea turtle cells. (a) Cellular respiration in lizard and (b) sea turtle cells across each treatment condition. (c) Cellular respiration in lizard and sea turtle cells exposed to hypoxia for 1 hour, followed by 1-hour reoxygenation. (d) Cellular respiration in lizard and sea turtle cells exposed to hypoxia for 24 hours, followed by 1-hour reoxygenation. Mitochondrial function: basal respiration, proton leak, maximal respiration, spare respiratory capacity, and ATP-linked respiration in lizard and sea turtle cells after (e) 1-hour hypoxia followed by 1-hour reoxygenation or (f) 24 hours hypoxia followed by 1-hour reoxygenation. Stars denote significant differences between species (p < 0.05, n=3).

### Preservation of the mitochondrial reticulum underlies the functional response to hypoxia/reoxygenation in sea turtle cells

We measured changes in mitochondrial morphology in cells exposed to hypoxia and hypoxia/reoxygenation to determine whether differences in the mitochondrial reticulum underlie the observed species-specific differences in primary carbon metabolites during hypoxia exposure and mitochondrial function during reoxygenation (**Figure 5a**). Intraspecific comparisons between hypoxia and hypoxia/reoxygenation showed significant differences in mitochondrial morphology for both species (p < 0.05). In lizard cells, network fragmentation increased after 1-hour hypoxia exposure (p = 0.0048), whereas mitochondrial footprint remained unchanged. In contrast, mitochondrial footprint (p = 0.020) and network branches decreased after 24-hour hypoxic exposure (p = 0.0028), suggesting further mitochondrial fragmentation.

**Figure 5.**
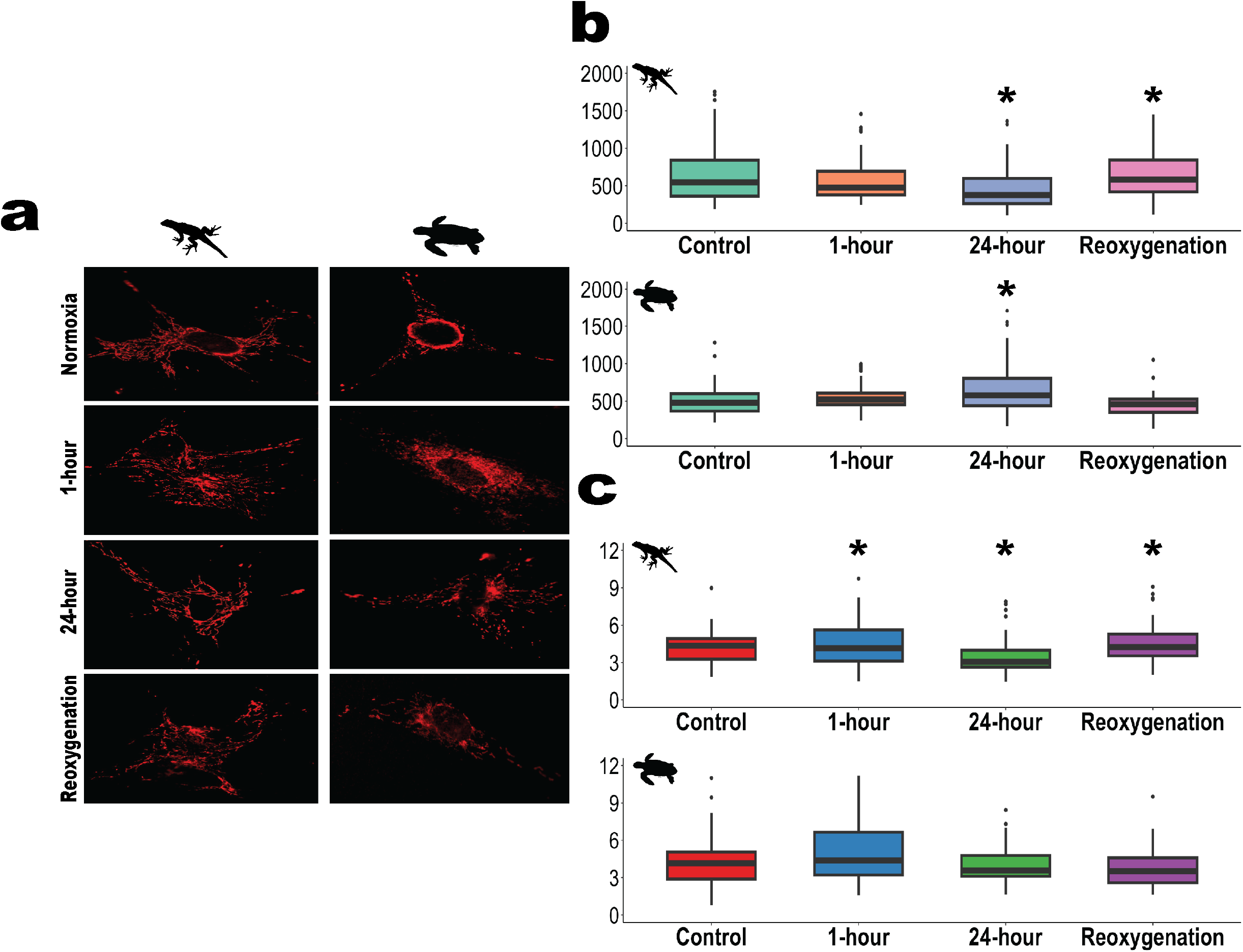
The mitochondrial reticulum is preserved during hypoxia exposure in sea turtle cells. (a) Cells from both species stained with MitoTracker red. (b) Mitochondrial footprint and (c) network branch counts in lizard and sea turtle cells under normoxia, 0.1% oxygen for 1 hour, 24 hours, and reoxygenation.

Mitochondrial morphology, however, began to recover after 24-hour hypoxia/reoxygenation (mitochondrial footprint: p = 0.020; network fragmentation: p = 0.0021). Thus, hypoxia exposure decreases mitochondrial footprint and increases network fragmentation in lizard cells. In contrast, hypoxia exposure for 24 hours increased mitochondrial footprint in sea turtle cells (p = 0.0213) and remained unchanged during reoxygenation (**Figure 5b, 5c**). Similarly, fragmentation of the mitochondrial reticulum upon hypoxia exposure was not observed in sea turtle cells. Hence, our studies suggest that sea turtle cells withstand long periods of hypoxia exposure by increasing their mitochondrial footprint, which likely drives the observed efficient recovery of cellular respiration during reoxygenation, in contrast to lizard cells. Altogether, these results suggest that during hypoxia, instead of switching to glycolysis like in lizard cells, sea turtle cells increase their mitochondrial footprint, promoting efficient oxygen use, while also supporting oxidative phosphorylation through circumventing the Crabtree Effect via galactose metabolism (**Figure 3d**). While reoxygenation damage might still occur in both species, as mitochondrial function is reduced (**Figure 4a-4d**), sea turtle cells can better maintain mitochondrial membrane integrity as more network fragmentation is seen in lizard cells. Furthermore, in contrast to lizard cells, sea turtle cells are able to restart mitochondrial function after extended hypoxia exposure (**Figure 5**).

## DISCUSSION

Freshwater turtles respond to hypoxia by undergoing metabolic rewiring while preserving mitochondrial function (Milton and Prentice, 2007; Fago, 2022; Sparks et al., 2022), but the metabolic adjustments sea turtles employ to withstand hypoxia at the cellular level had not been previously analyzed. Here, we compared the bioenergetic response to hypoxia exposure in primary cells derived from sea turtles and a non-diving reptile. Our results suggest that sea turtle cells use oxygen efficiently during hypoxia rather than relying on glycolysis. These changes are likely supported by the preservation of the mitochondrial reticulum and increased antioxidants that counteract reoxygenation-induced oxidative stress and allow efficient recovery of cellular respiration during reoxygenation.

### Glycolysis as a universal strategy to withstand hypoxia exposure?

HIF-1 reprograms metabolism, shifting carbon flux from oxidative phosphorylation to glycolysis during hypoxia exposure (Semenza, 2008; Baik and Jain, 2020). Our results suggest that lizard, but not sea turtle cells, exhibit a canonical metabolic switch to glycolysis upon exposure to hypoxia, despite pronounced HIF-1α stabilization in both species. Of note, although hypoxic sea turtle cells are likely not using the galactose pathway as an input for glycolysis, and metabolites in the glycolytic pathway are downregulated, we cannot conclude that glycolysis is not occurring without direct measurement. Hypoxia-tolerant vertebrates, such as the naked mole rat, rewire the glycolytic pathway, switching to fructose metabolism, which is converted to lactate in the heart and brain during oxygen-limiting conditions (Park et al., 2017). Similarly, anoxic freshwater turtles rely on glycogenolysis to maintain glucose levels during anoxia and balance low glycolytic ATP generation with low ATP requirements (Bundgaard et al., 2019a), suggesting that whole-body adjustments in carbohydrate metabolism play a crucial role in energy generation and use during oxygen-limiting conditions. In our experiments, we observed downregulation of gluconeogenesis in sea turtle cells, which might be intrinsic to using a cellular model. Nonetheless, our results show that while lizard cells become glycolytic during hypoxia, sea turtle cells do not appear to rely on the glycolytic pathway.

### Maintaining the mitochondrial reticulum during hypoxia exposure underlies efficient mitochondrial respiration during reoxygenation in sea turtle cells

Maintenance of the mitochondrial reticulum through mitochondrial dynamics (fission, fusion, and mitophagy) is a crucial process for oxidative phosphorylation, signal transduction (Picard and Shirihai, 2022), and the cellular stress response (Valente et al., 2017). We observed increased network branches, decreased mitochondrial footprint, increased abundance of glycolytic metabolites in lizard cells after 24-hour hypoxia exposure, and decreased cellular respiration during reoxygenation compared to sea turtle cells. In contrast, sea turtle cells exhibited an increased mitochondrial footprint, with no accumulation of glycolytic metabolites during hypoxia and sustained steady respiration upon reoxygenation. Our data are consistent with previous observations showing that freshwater turtles efficiently restart mitochondrial function after anoxia/reoxygenation (Fago, 2022). Thus, maintaining the mitochondrial reticulum appears crucial for supporting efficient oxygen use under limiting conditions and likely contributes to re-establishing mitochondrial function upon reoxygenation in hypoxia-tolerant reptiles. Consistent with our findings, previous work shows that mitochondrial integrity is maintained in anoxic turtle hearts (Bundgaard et al., 2019b), despite decreases in complex V activity (Gomez and Richards, 2018). Here, we show that mitochondrial integrity is maintained in sea turtle cells and that ATP-linked respiration increases significantly during reoxygenation after extended hypoxia. Proton leak signals many physiological functions, such as nutrient oxidation, mitochondrial oxidant production, glucose-stimulated insulin secretion, and decoupling carbon flux from ATP demand (Divakaruni and Brand, 2011). Our study shows lower overall cellular respiration but increased proton leak in lizard cells during reoxygenation after extended hypoxia exposure, suggesting higher oxidant production and reduced respiratory capacity. Taken together, our results show that after extended hypoxia, sea turtle cells can restore cellular respiration with minimal damage and that these adjustments result, at least in part, from the maintenance of the mitochondrial reticulum.

### Preparation for reoxygenation-induced oxidative stress in sea turtle cells

Animals adapted to drastic changes in oxygen availability have evolved diverse mechanisms to cope with reoxygenation-induced oxidative stress (Hermes-Lima and Zenteno-Savín, 2002). While freshwater turtles avoid oxidative damage by preventing hypoxia-induced succinate accumulation, which consequently limits mitochondrial superoxide generation upon reoxygenation, they also maintain high levels of antioxidants comparable to those in mammals, despite having much lower metabolic requirements (Storey, 1996; Hermes-Lima et al., 2015; Bundgaard et al., 2019a, 2023). Our results show significant enrichment of the glutamate pathway in sea turtle but not in lizard cells, suggesting that hypoxia promotes glutathione (GSH) synthesis (Krivoruchko and Storey, 2015). These results are consistent with findings on vascular endothelial cells derived from hypoxia-tolerant seal cells, where GSH is upregulated during hypoxia exposure (Allen et al., 2024). Together, these results support the idea that antioxidant upregulation during hypoxia is a conserved mechanism to cope with reoxygenation-induced oxidant generation in hypoxia-tolerant diving vertebrates (Zenteno-Savin et al., 2012). In addition, we also found that the antioxidant uric acid (Becker, 1993) is upregulated during both short and long-term hypoxia exposure in sea turtle cells. These data suggest that sea turtle cells sustain antioxidant protection during extended oxygen-limiting conditions, potentially preparing to cope with reoxygenation-induced oxidant generation.

In conclusion, we found species-specific changes in cellular respiration, metabolic rewiring, and mitochondrial morphology in response to hypoxia and reoxygenation exposure in primary cells derived from sea turtles and a non-diving reptile. Although hypoxia stabilizes HIF-1α in cells from both species, sea turtle cells upregulate antioxidant protection and are unlikely to rely on glycolytic metabolism during hypoxia. Furthermore, sea turtle cells can recover mitochondrial function during reoxygenation, and this functional response is likely underlined by the maintenance of the mitochondrial reticulum, which could promote efficient oxygen use during hypoxia-limiting conditions and efficient restart of cellular respiration upon reoxygenation. Understanding the cellular changes driving natural protection against reoxygenation-induced damage could provide translational information for treating ischemic injuries characterized by drastic changes in oxygen availability.

## ACKNOWLEDGMENTS

We thank Noemi Bautista, Johnny Hoang, Yvette Javier, Mohamed Moustafa, Joseph Nguyen, Amber Singh, and Harshmeet Singh for their help with lizard collection and husbandry. Federico Kong-Gonzalez, Eqlima Tahiry and Dua Shoaib helped with wet lab analyses. BGA was supported by Ford Foundation predoctoral and dissertation fellowships. JPV-M is supported by NIGMS grant R35GM146951. Research funded by UC Berkeley.

## FIGURE LEGENDS

**Supplementary Figure 1.**Enrichment analyses of significant metabolites mapped to pathways in the SMPDB database, with a minimum of three entries required per pathway. Pathways with significant enrichment are highlighted in green.

