## Supplementary figures and images for "Hypoxia exposure fine-tunes mitochondrial function in sea turtle cells"

### Supplemental Figure 1

#### Supplementary Figure 1.

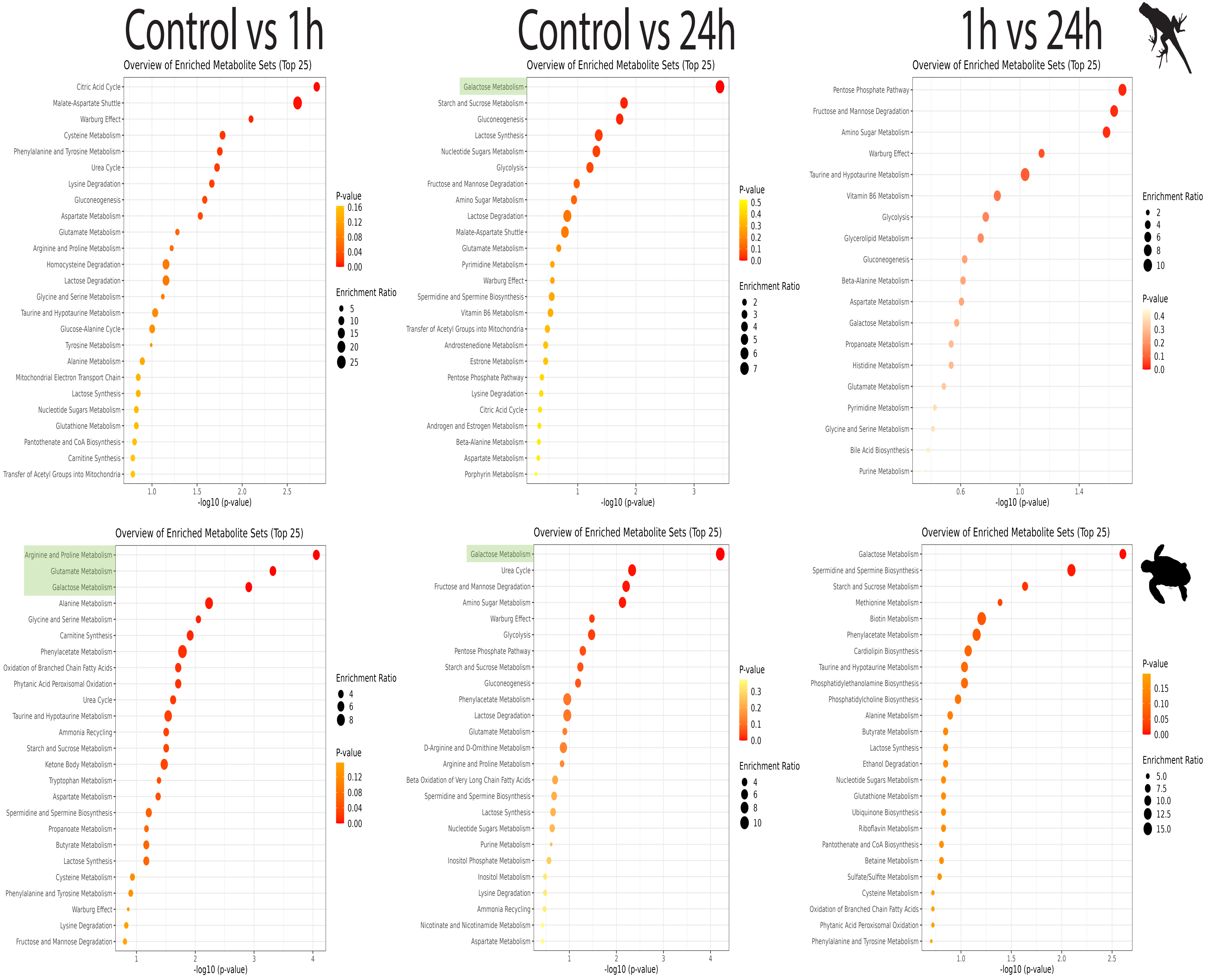
